# Fast clustering and cell-type annotation of scATAC data using pre-trained embeddings

**DOI:** 10.1101/2023.08.01.551452

**Authors:** Nathan J. LeRoy, Jason P. Smith, Guangtao Zheng, Julia Rymuza, Erfaneh Gharavi, Donald E. Brown, Aidong Zhang, Nathan C. Sheffield

## Abstract

**Motivation:** Data from the single-cell assay for transposase-accessible chromatin using sequencing (scATAC-seq) is now widely available. One major computational challenge is dealing with high dimensionality and inherent sparsity, which is typically addressed by producing lower-dimensional representations of single cells for downstream clustering tasks. Current approaches produce such individual cell embeddings directly through a one-step learning process. Here, we propose an alternative approach by building embedding models pre-trained on reference data. We argue that this provides a more flexible analysis workflow that also has computational performance advantages through transfer learning.

**Results:** We implemented our approach in scEmbed, an unsupervised machine learning framework that learns low-dimensional embeddings of genomic regulatory regions to represent and analyze scATAC-seq data. scEmbed performs well in terms of clustering ability and has the key advantage of learning patterns of region co-occurrence that can be transferred to other, unseen datasets. Moreover, pre-trained models on reference data can be exploited to build fast and accurate cell-type annotation systems without the need for other data modalities. scEmbed is implemented in Python and it is available to download from GitHub. We also make our pre-trained models available on huggingface for public use.

**Availability:** scEmbed is open source and available at https://github.com/databio/geniml. Pre-trained models from this work can be obtained on huggingface: https://huggingface.co/databio.

## Introduction

Data from the single-cell assay for transposase-accessible chromatin using sequencing (scATAC-seq) can interrogate complex regulatory networks at the single-cell level, elucidating the cellular mechanisms that drive cell-to-cell heterogeneity. The power of scATAC-seq data has motivated the development of new computational approaches for its analysis [1–12]. Despite these advances, scATAC-seq analysis continues to face two key challenges: the 1) high dimensionality and 2) inherent sparsity of the data [13, 14].

scATAC-seq analysis often includes two critical tasks: 1) dimensionality reduction followed by clustering and 2) cell-type annotation of cell clusters. For the dimensionality reduction task, numerous methods have been developed, such as SCALE and scBasset, which use variational autoencoders and convolutional neural networks to learn low-dimensional representations of single cells for downstream clustering tasks [1, 2]. Other methods include ChromVAR, cisTopic, SnapATAC, and ArchR, which leverage latent semantic indexing (LSI) and topic modeling to cluster individual cells [3–5, 9]. These methods usually require complex processing pipelines and large compute power. The second task, cell-type annotation, is less well served, with most current methods simply repurposing cell-type annotation tools from scRNA-seq [15]. Methods developed for scATAC are few and suffer notable limitations. First, they mainly take a cross-modality approach, integrating data from reference scRNA-seq sets, so they are limited in relying on a secondary data modality. Finding an appropriate secondary dataset to complement the unlabeled set can be difficult [16]. Second, many supervised methods require model training to predict cell types from a fixed output. This can make the discovery of novel cell types a challenge [12]. Finally, these methods are notoriously compute-intensive [17], a limitation that has grown more problematic as atlas-level datasets have emerged.

Here, we address these challenges with an alternative approach to scATAC-seq dimensionality reduction and cell-type annotation using pre-trained embedding models. Our method improves both dimensionality reduction and cell-type annotation by significantly reducing the computational time and complexity of these workflows with the added benefit of leveraging information from highquality reference data sets. Instead of analyzing datasets end to end, we use unsupervised learning to model the patterns of regulatory region co-occurrence in reference datasets, and then transfer this knowledge to new, unseen data. While other transfer learning methods do exist, nearly all focus on the integration of scRNA-seq data with scATAC-seq [18–20]. Moreoever, methods that do focus on transfer learning for scATAC-seq data [21] leave much to be desired. First, they lack the ability to utilize publicaly available pre-trained models; second, they have yet to be used for cell-type annotation; and third, they lack a software framework to facilitate easy analysis. Such a framework could drastically speed up analysis. To that end, we designed our method to remedy these shortcomings. We implemented this method in scEmbed, an un-supervised machine-learning method that learns low-dimensional representations of genomic regions from scATAC-seq datasets.

We first show that scEmbed performs well for dimensionality reduction and clustering while maintaining robustness to data loss. Moreover, by leveraging models pre-trained on reference data, scEmbed drastically reduces the time and complexity of scATAC-seq analysis. Finally, we build a cell-type annotation system by exploiting the learned embeddings produced by pre-trained embedding models without needing any external data modalities. Our system can accurately annotate unseen data in seconds using pre-trained reference models. scEmbed takes a new approach to scATAC-seq analysis by focusing on ATAC-seq data alone, by building high-quality embeddings of genomic regions en route to single-cells, which offers flexibility and speed for a wide range of downstream tasks.

## Results

### scEmbed architecture

scEmbed adapts our previous work, Region2Vec [22], to single cells. The model is a modified unsupervised word2vec [23] model that learns to predict genomic region co-accessibility (Fig. 1A). Briefly, scEmbed treats each cell as a document and its accessible regions as words (see Methods). Context is simulated by shuffling these regions in each document (Fig. 1B). After training, cell embeddings are constructed by averaging region vectors for each cell, which are then used for tasks like clustering, analysis, or transfer learning (Fig. 1C).

**Figure 1.**
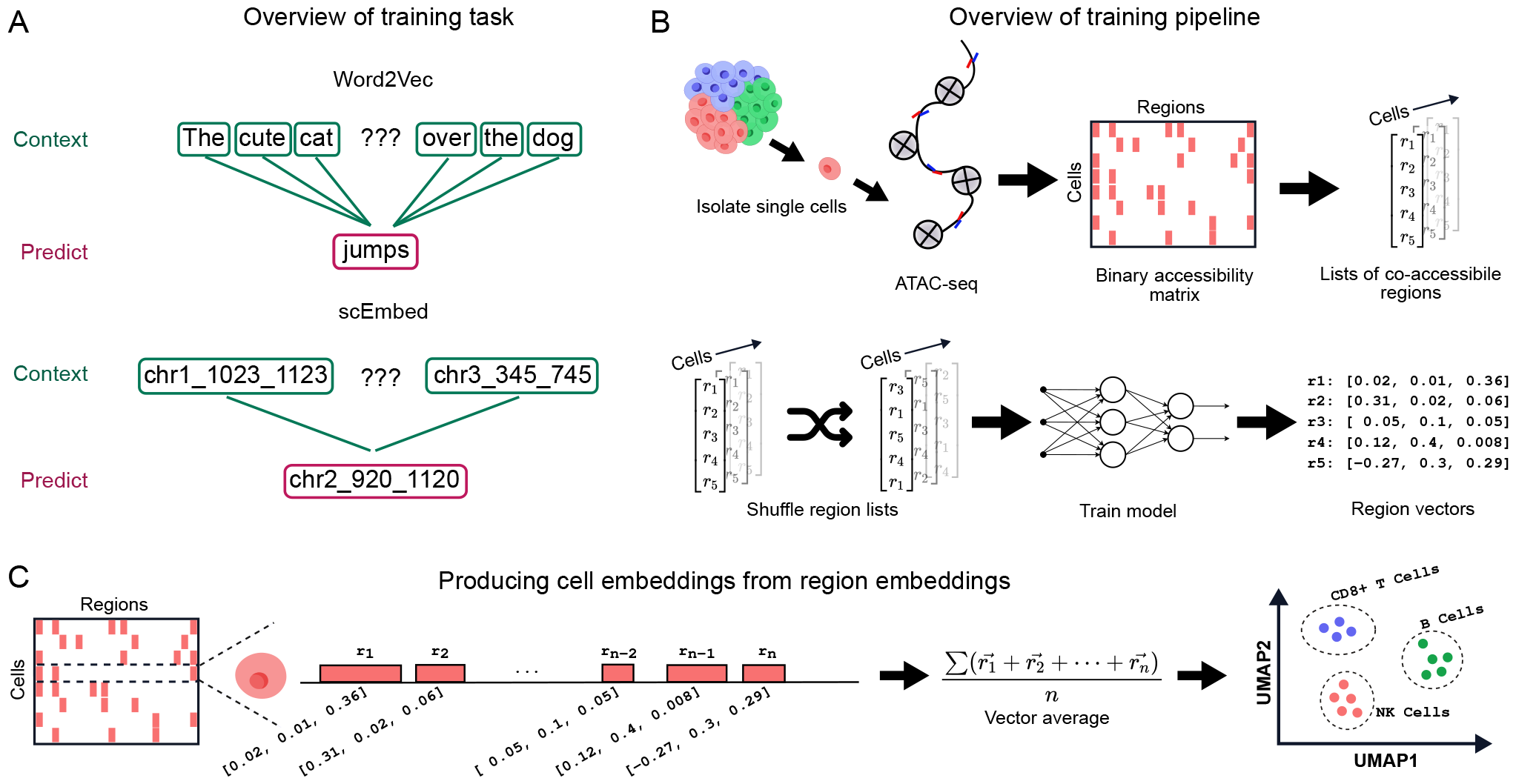
Overview of the scEmbed model. **A**. scEmbed leverages Word2Vec as its model. Word2Vec learns to predict words given a semantic context. Similarly, scEmbed learns to predict genomic regions, given a genomic context. This is unsupervised, and uses the patterns of genomic region co-occurrence to learn representations of individual regions. **B**. Overview of the scEmbed learning process, starting with scATAC-seq data. **C**. Once region embeddings are learned, they can be used to construct cell embeddings by averaging the embeddings of regions accessible in each cell. We use cell embeddings for downstream tasks of clustering and cell-type prediction.

### scEmbed model validation

To validate scEmbed, we followed an earlier approach [13] to benchmark it on clustering tasks using published reference scATAC data from hematopoietic cells [24] (Fig. 2A). We trained scEmbed for 100 epochs (Fig. S1) then used the resulting region embeddings to construct cell embeddings. Visually, scEmbed clusters cells of the same type (Fig. 2B). Following previous benchmarking procedures, we clustered the cell embeddings with three clustering methods: K-means, hierarchical clustering (HC), and Louvain clustering. The clusters were then compared to ground truth labels using three metrics: adjusted rand index (ARI), the adjusted mutual information score (AMI), and the homogeneity score (see Methods). scEmbed performs similar to the best-performing scATACseq methods, including SCALE, scBasset, cisTopic, and SnapATAC (Fig. 2C). It does so with almost no preprocessing of the data and a completely unsupervised learning workflow. In addition to the Buenrostro2018 dataset, we also benchmarked scEmbed on another, more recent and comprehensive scATAC-seq dataset from Luecken *et. al* [25]. Again, comparing clusters to ground truth labels, scEmbed performs well (Fig. S2).

**Figure 2.**
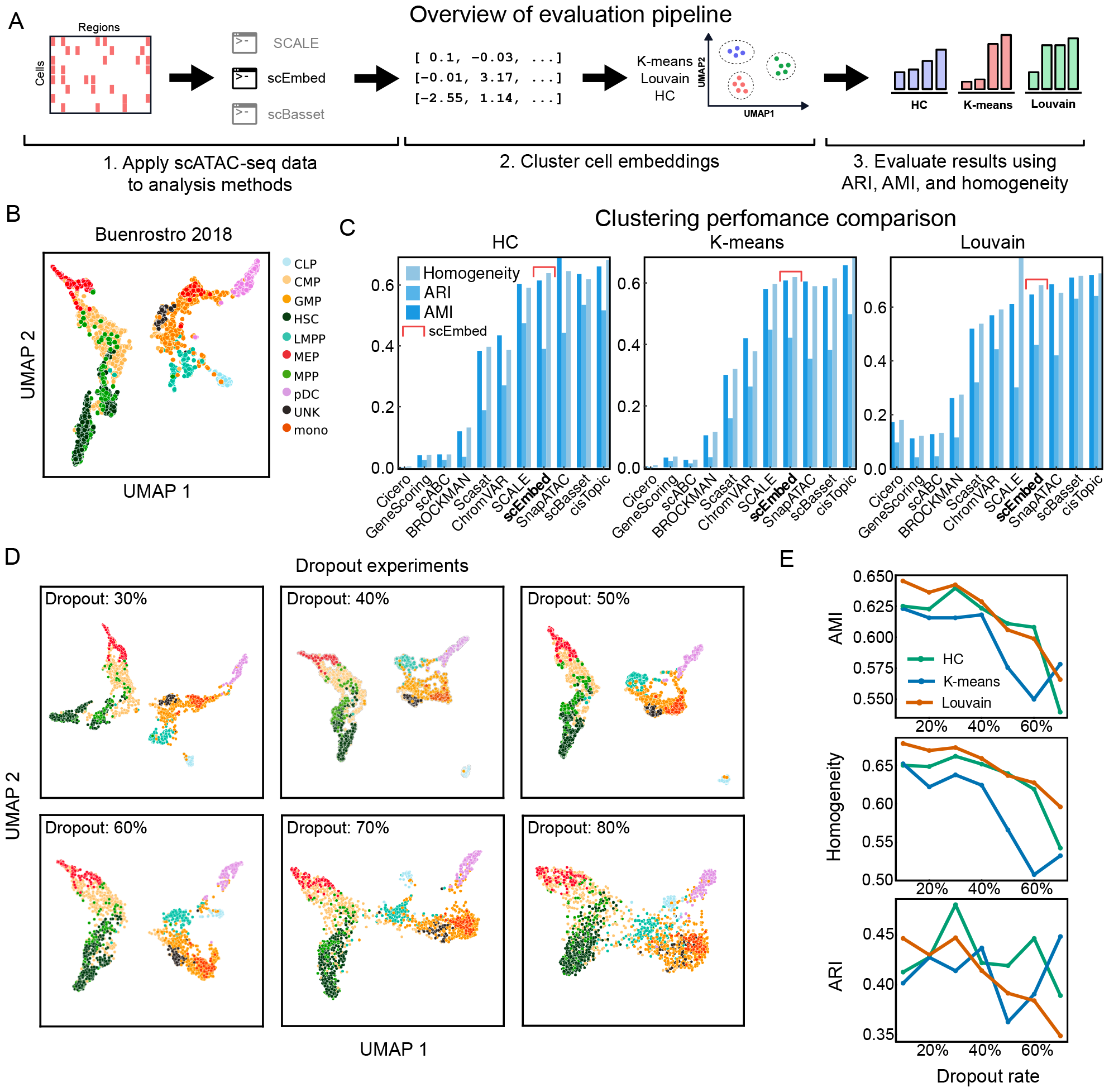
scEmbed benchmarks competitively with existing approaches. **A**. Diagram showing 3 steps of the benchmarking process. **B**. UMAP of scEmbed cell embeddings. **C**. Benchmark results of 3 clustering methods: Hierarchical clustering (HC), K-means, and Louvain. Results were evaluated using three metrics: adjusted mutual information (AMI), adjusted rand index (ARI), and homogeneity. **D**. UMAP plots showing clusters of scEmbed cell embeddings following data loss. **E**. Line plots showing the change in three clustering metrics (ARI, AMI, and Homogeneity) as a function of dropout rate. scEmbed accurately clusters single cells up to nearly 80% data loss.

### scEmbed is robust to data loss

We were curious to explore the potential of scEmbed for transfer learning tasks. One common challenge in transfer learning is handling the nuances and disparities between the original training data and new data on which inference is performed. These differences can sometimes lead to a perceived loss of information or data inconsistencies. As such, we sought to evaluate its ability to cluster data with increasing levels of information loss. Our experiments **began** with an already sparse cell-feature matrix. From there, we dropped out values until the matrix was **even more** sparse. To test scEmbed’s robustness to missing data, we trained the model on these datasets of increasing sparsity. Starting with the Buenrostro2018 dataset (2.8% non-zero) [24], we randomly dropped non-zero values in the binary accessibility matrix until approximately 80% of the non-zero data was lost. A dropout rate of 80% resulted in a matrix that was 0.5% non-zero (Methods, Fig. S3). Even at a dropout rate of 80%, scEmbed was able to visually cluster cells of the same type (Fig. 2D). To quantify this, we computed three scores for each dropout dataset: 1) Adjusted Rand Index (ARI), 2) Adjusted Mutual Information (AMI), and 3) Homogeneity scores. We found that scEmbed retained clustering accuracy even when faced with substantial data loss (Fig. 2E). This is comparable to other scATAC clustering methods which are robust to data loss [13]. These findings confirm that scEmbed can learn rich biological knowledge, even for the most sparse datasets. The ability to handle sparseness is a critical characteristic of scATAC-seq analysis, and particularly so for scEmbed, which can be used to transfer information from existing models. Knowledge is transfered through region overlap analysis which is an imperfect process. Therefore scEmbed should learn even when data is incomplete. We describe this next.

### scEmbed transfers knowledge of genomic region cooccurrence to unseen datasets

A key innovation in scEmbed is that it uses a two-step training process, rather than the common single step approach (Fig. 3A). In the first step, scEmbed learns embeddings of genomic regions rather than cells. In the second step, the region embeddings are used to build cell embeddings. An advantage of this two-step approach is that the region embeddings can be used to build cell embeddings for new datasets. This transfer approach allows scEmbed to take advantage of pre-trained reference models. We call this “projection” because we “project” new data into the latent space of the original dataset, creating cell embeddings for new data using a pre-trained model. Projection occurs in three steps: First, we train a model on reference data to produce region embeddings for each region in the reference consensus region set. Second, we take a new single-cell dataset and map the regions to the reference consensus region set using region overlaps (Fig. 3B, Methods). This represents each single cell in the new dataset using the set of regions from the reference dataset, for which we also have region embeddings from the reference model. Finally, we compute the average of all region embeddings for each cell in the new dataset (Fig. 3C). This approach leverages the information from a larger atlas of accessibility data to analyze a new dataset. In fact, the original training data need not come from scATAC-seq at all. Using this approach, a model trained with bulk ATAC-seq could similarly be used to project scATAC-seq data. This provides an enormous advantage by utilizing the patterns of region co-occurrence from the vast volume of publicly available region set data to inform cell embeddings of single-cell data.

**Figure 3.**
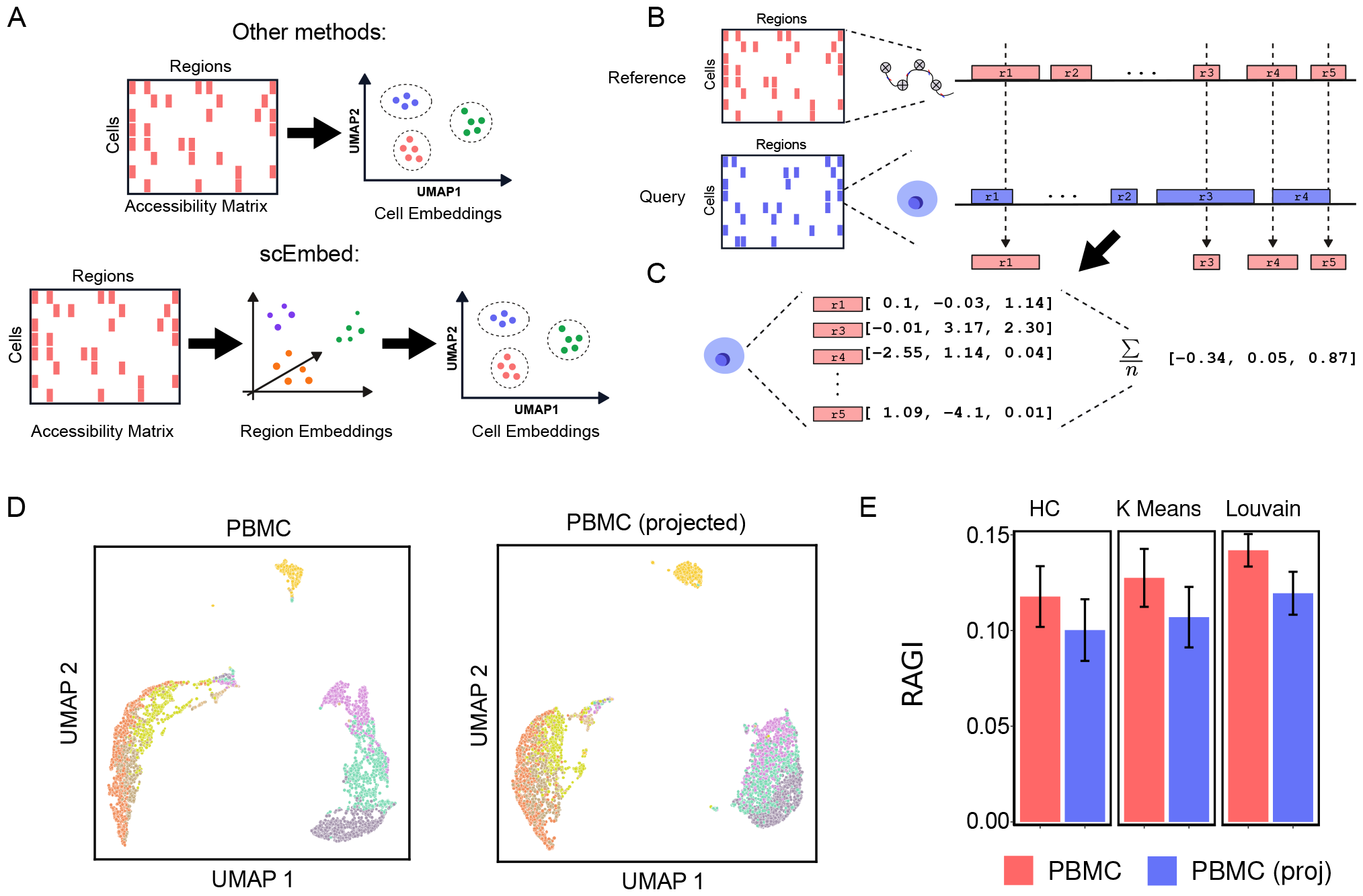
scEmbed enables knowledge transfer to unseen datasets. Transfer learning with scEmbed occurs in three steps. **A**. Diagram showing the high-level workflow of other scATAC-seq methods (top) compared to scEmbed (bottom) **B**. Diagram of the overlap analysis procedure. Using interval overlap analysis, a new cell from a new dataset (blue) can be cast in the feature space of the data used for the pre-trained model (red). **C**. Diagram showing the computation of embeddings for new, unseen data. This is achieved using basic average pooling of region embeddings. **D**. UMAP plots of both projected (right) and unprojected (left) datasets. The plots show nearly identical clustering of embeddings learned from the original dataset versus projection. **E**. RAGI score plots for both the original dataset embeddings and projected cell embeddings using a repeated sampling strategy. RAGI scores are computed as the mean RAGI score across all samples for three clustering methods: Hierarchical clustering, K-means, and Louvain. Error bars indicate one standard deviation away from the mean.

### Projected cell embeddings cluster cells accurately using pretrained models

We next assessed this projection process by asking whether scEmbed could cluster a new dataset based entirely on a pre-trained model. First, we trained a model on the original Buenrostro2018 dataset [24]; second, we took a new dataset, 10X genomics 5k Peripheral blood mononuclear cells (PBMCs) from a healthy donor, and projected each cell into the original space. We used these single-cell embeddings directly for UMAP visualization and clustering analysis. To assess the quality of the projection, we assumed that 8 distinct cell populations existed [13] and took advantage of marker gene analysis to assign labels to each cell. We use the Residual Average Gini Index (RAGI) score to evaluate the clustering ability of scEmbed [13] (Methods).

We found that the projected-cell analysis showed no marked differences in clustering proficiency when compared to the embeddings produced by conventional model training. The UMAP plots were visually similar (Fig. 3D) indicating similar clustering performance. To further explore the difference, we next evaluated clustering performance using a repeated subsampling strategy (Fig. S4) that consisted of four steps: 1) train a new model on the PBMC data alone; 2) repeatedly subsample 1,000 cells and compute their embeddings using the new model and the Buenrostro2018 model using projection; 3) cluster the cells using three strategies (HC, Kmeans, and Louvain); and 4) compute the RAGI score with these 1,000 subsampled cells. The scores were then averaged across all subsamples. Our results showed that the RAGI score between the original and projected datasets did not differ significantly, indicating similar clustering performance (Fig. 3E).

### Pre-trained models from reference datasets can be used to annotate cell clusters

Given the promising results of our initial experiments with pretrained models, we next sought a way to visualize the projected cells. Furthermore, we reasoned that this approach could be used to annotate cell clusters for the projected cells, allowing us to borrow annotation information from the reference model. To build such a system, we distinguish between three data flows that can occur with scEmbed (Fig. 4A). The first data flow is the *no projection* flow. This is the standard workflow of training a new scEmbed model on some input data and visualizing the resulting embeddings by fitting a UMAP model to reduce the dimensionality to 2. The second data flow is the *embedding-only projection workflow* (E-projection). In this data flow, new data is embedded using a pre-trained model trained on reference data, as described above. These embeddings may then be visualized by fitting a UMAP model to reduce dimensionality to 2 dimensions. The final data flow, and the novel innovation that accomplishes our goal of reference-based visualization, is the *embedding and visualization workflow* (EV-projection; Fig. 4A). In this data flow, new data is first embedded using a pre-trained model on reference data, as in E-projection; then, these embeddings are further projected through a UMAP model that was *fit on the reference data embeddings*, rather than newly fit. With the EV-projection flow, plotting the 2-dimensional cell representations on top of the reference data UMAP plot allows one to visualize where in the original embedding space the new data ended up (Fig. 4B). This is possible because the EV-projection re-uses the same topology from the UMAP model fit to the reference data (Fig. 4C).

**Figure 4.**
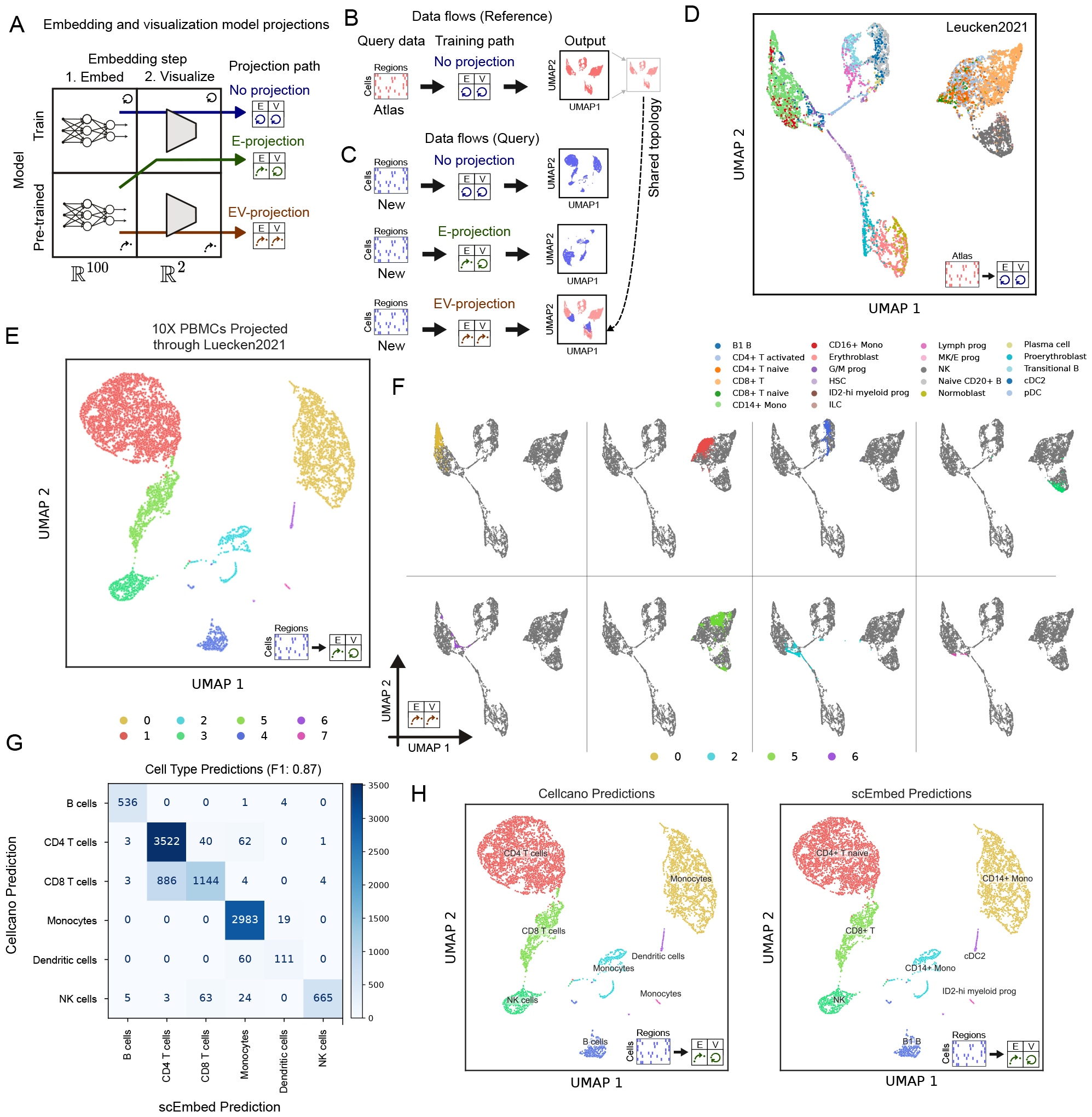
Pre-trained embedding models can be exploited for cell-type annotation tasks. **A**. Diagram showing scEmbed’s three projection paths. **B**. Overview of the standard “no-projection” data flow. **C**. Overview of three data flows for new data. EV-projection places the new data in the same latent space as the reference data. **D**. UMAP plot of the reference data embeddings built using the standard workflow. **E**. UMAP plot of the new PBMC data with E-projection. **F**. Plots showing the EV-projection data flow applied to the new PBMC dataset. Grey cells represent the reference topology; colored cells are projected new PBMC data. Separate plots depict individual clusters for visual clarity. **G**. Confusion matrix of scEmbed classification results compared to Cellcano. **H**. UMAP plots showing the cell labels assigned by Cellcano (left) and the cell type labels assigned by scEmbed (right).

To demonstrate this approach for reference-based visualization and annotation of new data, we built a reference model using the Luekcen2021 multi-omic dataset [25], a first-of-its-kind multimodal benchmark dataset of 120,000 single cells from the human bone marrow of 10 diverse donors measured with two commercially available multi-modal technologies. Using scEmbed and the “no projection” data flow, we trained a model and clustered the resulting embeddings (Fig. 4D). This model served as the reference for all downstream experiments with a new PBMC dataset from 10X genomics. Using E-projection, scEmbed creates visually distinct clusters of single cells (Fig. 4E, Fig. S5). To visualize these embeddings in the context of the original embedding topology, we employ EV-projection. Each identified cluster from E-projection aggregates to a distinct location in the original UMAP embedding topology of the Luecken2021 model (Fig. 4F).

Confident our pre-trained Luecken2021 embedding model was distinctly clustering the new PBMC dataset, we sought to assign cell-type labels to each cluster. We leveraged Cellcano, a new scATAC-seq cell-annotation method, to assign ground-truth labels to each cluster of the E-projected PBMC embeddings for evaluation of our method [12]. Our cell-type annotation system was limited by the cell types annotated in Luecken2021; as such we mapped each scEmbed prediction class to a corresponding Cellcano class for comparison (Table S1) after following their annotation procedure (Methods, Fig. S6). Using a simple k-nearest-neighbor (KNN) classification algorithm (see Methods), scEmbed was highly consistent with the Cellcano labels (F1=0.87, Fig. 4G). However, without class mapping, scEmbed offers higher specificity of cluster identity and even identifies a cluster of ID2-hi myeloid progenitor cells not found with Cellcano (Fig. 4H). Moreover, this workflow enables researchers to quickly try new models trained on many different cell types and rapidly discover cell types in their data. Using our projection system, researchers can avoid training a new model each time they want to use a new reference data set, which is a common approach in many modern cell-type annotation systems [7, 12, 26]. The entire process of dimensionality reduction, clustering, and annotation took less than 10 minutes on a laptop, and we observed similar time savings across several models (Fig. S7). Thus, we conclude that EV-projection is a promising approach for fast visualization and annotation of new data.

## Discussion

In this work, we demonstrate the robustness and versatility of scEmbed, a new tool for the analysis of scATAC-seq data. scEmbed differs from existing methods in that instead of learning embeddings of individual cells directly, it first learns embeddings of genomic regulatory regions and then uses these to compute cell embeddings. We demonstrate how this approach allows scEmbed to use pre-trained genomic region embedding models to effectively cluster data not seen by the model. Our evaluation of scEmbed against existing scATAC-seq methodologies demonstrates its efficacy and competitiveness, even with a relatively simple network architecture. scEmbed performs well, even when faced with severe data loss. The standout feature of scEmbed is its capability to repurpose learned region embeddings for downstream analysis tasks. This approach provides flexibility and efficiency, setting it apart from other currently available tools.

By exploiting region overlaps and applying previously learned region embeddings, we have formulated a novel method for representing unseen scATAC-seq data within the latent space of the original training data. This process, termed “projection,” yielded superb clustering of cells, showing no significant decrease in performance compared to models trained entirely on the new dataset. This performance underscores the potential of scEmbed in the context of ATAC-seq transfer learning tasks and opens exciting possibilities for future research. Moreoever, we emphasize the novelty of scEmbed in its ability to engage in transfer learning without the need for another data modality like scRNA-seq, which is overwhlemingly required by current methods. Future studies will explore the ability of our model to learn and extract overarching regulatory patterns from publicly available data. This learning, coupled with the inherent transferability of scEmbed, will empower researchers to fine-tune the models for specific downstream tasks, enabling gains in performance, efficiency, and flexibility.

Finally, we leveraged embeddings computed by scEmbed and its pre-trained models to build a novel cell-type annotation system. Our method is consistent with current scATAC-seq cell type annotation implementations with the added advantage of requiring no external data modalities. Furthermore, by exploiting pre-trained models and pre-computed cell embeddings from reference datasets, the scEmbed annotation system can easily scale to millions of cells and uses only a fraction of the compute time. This is because utilization of a pre-trained model, also known as projection, requires only interval overlap analysis to map the new data into the feature space the model was trained on (Fig. S7). We’ve made the pre-trained models used in this paper available for download and use on huggingface. To facilitate model sharing and usability even further [27], we’ve built software packages to easily download and use these models within Python. Moreover, these same packages can be used to train new models or fine-tune public ones on custom datasets. We hope that these resources will enable researchers to leverage the power of scEmbed for their own research.

The integration of unsupervised learning with transfer learning may offer new directions for other bioinformatics tasks that are similarly burdened by the challenges of high-dimensionality and data sparsity. Furthermore, the deployment of pre-trained models for reference datasets may inspire novel methodologies for efficient and accurate cell-type annotation systems across different data modalities. In the future, the pre-training approach of scEmbed could be adapted for use with cross-modality methods that span data types [28]. In conclusion, scEmbed’s ability to distill meaningful representations from vast, complex scATAC-seq datasets, and repurpose this knowledge for rapid and accurate analysis of new datasets, has great potential. This work is a step towards developing more efficient, scalable, and flexible tools for genomic data analysis. The opportunities unlocked by scEmbed for research and clinical application promise exciting advancements in the comprehension of cellular heterogeneity and the intricate regulatory networks that drive it.

## Methods

### Data and data processing

#### Overview of datasets

##### Luecken2021

The Luecken2021 dataset is a multimodal singlecell benchmarking dataset [25]. The data is a first-of-its-kind multimodal benchmark dataset of 120,000 single cells from the human bone marrow of 10 diverse donors measured with two commercially-available multi-modal technologies: nuclear GEX with joint ATAC, and cellular GEX with joint ADT profiles. The data was retrieved from the gene expression omnibus (GEO) using the GEO accession GSE194122.

##### Buenrostro2018

The Buenrostro2018 dataset consists of singlecell chromatin accessibility profiles across 10 populations of immunophenotypically defined human hematopoietic cell types [24]. Deduplicated single-cell bam files along with a consensus peak set were provided by Chen *et. al*. [13]. For datasets where consensus peaks don’t exist, methods that create such sets from raw data could be used as a pre-processing step [29]. Using `bedtools` [30], region overlaps with the consensus peak set were computed for each bam file at a minimum overlap of 1bp. Using the `-c` flag, the number of overlaps with each region in the consensus peak set was calculated. Overlap count files were subsequently converted into a cell by peak binary accessibility matrix formatted as a comma-separated-value file (`csv`). Finally, the binary accessibility `csv` was converted into a `scanpy AnnData` object using the `scanpy.read csv` API. This was used as input to the scEmbed model.

##### 5k PBMC

The PBMC dataset comes from 10X genomics and consists of peripheral blood mononuclear cells (PBMCs) from a healthy donor. Three files were downloaded directly from the 10X genomics website: 1) the sparse peak matrix in `.mtx` format, 2) the cell barcode labels in `tsv` format, and 3) the consensus peak set in `bed` format. Using Python, along with `pandas` and `scanpy`, these files were processed into a `scanpy AnnData` object. This was used as input to the scEmbed model.

##### Synthetic Bone Marrow

The synthetic bone marrow dataset was described and provided by *Chen et. al*.[13]. The binary accessibility matrix was downloaded directly from the Pinello Lab’s GitHub as a `.rds` file. Using R, this file was read, parsed, and exported as a `csv`. Like the previous two datasets, this `csv` was processed into a `scanpy AnnData` object using `pandas` and `scanpy`.

### Architecture of scEmbed

#### Word2Vec model

We used the `gensim` implementation of Word2Vec as the core model for scEmbed. Word2Vec has many configurable hyperparameters [31], including context window size, embedding size, learning rate scheduling, and number of epochs. All experiments were conducted with a fixed set of hyperparameters. We used defaults for scEmbed, informed by experiments on Region2Vec optimization [32]. Specifically, we use a window size of 5 and an embedding dimension of 100. We also use 100 epochs for all experiments unless otherwise noted. We adopt an exponential learning rate schedule with a decay rate of 0.95.

#### Presenting cells as documents to Word2Vec

Word2Vec takes lists of words as input. To this end, we designed a way to convert a binary accessibility matrix into list of words that are compatible with Word2Vec’s acceptable input corpus format. scEmbed expects binary accessibility matrices with cells as rows and regions as columns. Briefly, we treat each cell as a sentence and each co-accessible region in the cell as a word. Using this convention, we generate input for Word2Vec in three steps. First, we take an individual cell and identify each region where it shows signal. We define signal as anything greater than zero. Second, we take the corresponding region that shows signal and convert it into a word by concatenating the chromosome, start, and end values with an underscore (`chr start end`). This process is completed for each region with signal in the cell, and a list of “words” is constructed. Finally, this list is shuffled repeatedly to simulate context and used directly as input to Word2Vec. Shuffling is necessary since co-accessible regions have no inherent order, and Word2Vec learns by context. We repeat this process for each cell in the accessibility matrix.

### Clustering

We use three clustering algorithms: Hierarchical clustering (HC), k-means clustering, and Louvain clustering. For HC and k-means, we use the `scikit-learn` implementations. When ground-truth labels were known for a particular dataset, we used the number of unique labels to set the number of clusters to generate. Otherwise, we used prior knowledge to estimate the number of unique cell populations we would expect to find. For Louvain clustering, we use the `scanpy` implementation. Louvain is agnostic to a specified number of clusters. As such, we iteratively applied clustering to datasets while slowly increasing the resolution value from 0 to 3. With each iteration, the number of clusters was stored in a list along with the corresponding resolution. Once complete, we employed binary search on the list to identify the resolution that gave us the desired number of clusters. This value was used to generate the final clustering solution.

### Visualization

We used uniform manifold approximation and projection (UMAP) to visualize single-cell embeddings [33]. We used the `umap-learn` Python package and specified two dimensions for each visualization. In addition, a random state of 42 was set for visualization workflows. All other parameters were set to package defaults.

### Clustering evaluation

#### Evaluation metric

There are two scenarios for which we can evaluate clustering: known ground-truth labels and unknown ground-truth labels. The synthetic bone marrow and Buenrostro2018 datasets constitute the known ground-truth label scenarios while the PBMC data constitutes the unknown ground-truth label scenario. When groundtruth labels are known, we can employ three different scores: The adjusted rand index (ARI), the adjusted mutual info score (AMI), and the homogeneity score. When ground-truth labels are not known, we use the Residual Average Gini Index (RAGI) [13].

#### Adjusted Rand Index

The ARI is a metric for evaluating the similarity between two data clusterings. This is achieved by counting pairs that are assigned to the same cluster label. Mathematically, it is computed by:

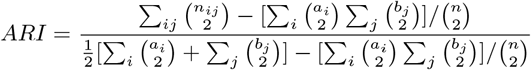

where *n*_*ij*_, *a*_*i*_, *b*_*j*_ are diagonal values, row sums, and column sums respectively from the contingency table that describes the frequency distribution of the cluster labels from ground-truth and predicted clusterings.

We use the `adjusted rand score` function from the `scikit-learn` python package.

#### Adjusted Mutual Info Score

The AMI, intuitively, is a measure of the amount of information that two clusterings share. It’s used to evaluate how well two clusterings agree with each other [34]. We compute AMI through the `scikit-learn` package using the `adjusted mutual info score` function.

#### Homogeneity Score

The homogeneity score is an entropy-based external cluster evaluation metric that measures how far from perfect an incorrect clustering solution is [35]. We employ the `scikit-learn homogeneity score` function to measure this metric for each dataset.

#### Residual Average Gini Index

When ground truth labels are unknown, all aforementioned evaluation metrics are no longer applicable. As such, we need a measure that can still evaluate dataset clustering based on what one would expect, given some sort of prior knowledge about the system. For this, we employ a similar strategy described by *Chen et. al*. called the Residual Average Gini Index (RAGI). Briefly, the RAGI score compares the accessibility of housekeeping genes with previously characterized marker genes [36]. RAGI measures the average residual specificity of a clustering solution with respect to marker genes, suggesting that a good clustering solution should have clusters enriched for different marker genes and these genes should be highly accessible in only a few clusters, compared to the less informative housekeeping genes.

### Data corruption and dropout experiments

scATAC datasets are notoriously sparse. To that end, we wanted to investigate scEmbed’s ability to handle increasingly sparse matrices that go beyond even a standard scATAC-seq dataset. To simulate this increasing information loss, we began with a standard scATAC-seq binary accessibility matrix and then iteratively and randomly dropped non-zero values in the matrix down to zero at increasing rates. Following a similar approach described by *Xiong et. al*. to evaluate SCALE [1], we randomly dropped out non-zero values at increments of 10% from 0.1 to 0.8. This resulted in increasingly more sparse feature matrices (Fig. S3). These matrices have been made more sparse than the original, already-sparse scATAC binary accessibility matrix. Using `choice` from the `numpy` python library, we changed all non-zero values in the feature matrix to 0 with a probability of Dropout Rate. The resulting matrix was saved and used for downstream analysis.

### Transfer learning and data projection

#### E-projection

The E-projection method enables the transfer of learned knowledge from a pre-trained model to unseen datasets. It translates a new binary accessibility matrix into the space of the original dataset. For each cell in the new dataset, this projection occurs in three stages:

##### Region Overlap Computation

We first calculate region overlaps between the accessible regions in the new cell and the consensus peak set from the original dataset. This is achieved using interval trees, a data structure that facilitates efficient discovery of all intervals that overlap with any given interval or point. We use the `intervallist` package in python for these computations. To increase speed and performance for on-disk datasets, the augmented interval list (AIList) datastrecture could be used [37]. Regions with signal that overlap are mapped onto the original dataset’s consensus peak set.

##### Word-Key Conversion

Subsequently, each of the overlapping regions is transformed into its corresponding word-key. This is accomplished by joining the values of the chromosome, start, and end with an underscore (`chr start end`).

##### Embedding Transformation and Averaging

Finally, these word-keys are translated into their corresponding embeddings. Region embeddings are obtained by passing the regions corresponding onehot vector through a single embedding layer. We denote a one-hot encoded vector for a given region as *O*_*r*_, where *r* represents the specific region. We then denote the embedding weight matrix as *W*_*e*_, where each column corresponds to the embedding of a particular region in embedding space. Given these symbols, the operation to translate a one-hot encoded vector into its corresponding embedding can be represented as: *E*_*r*_ = *W*_*e*_ *· O*_*r*_ . An average is then calculated across these embeddings to derive a whole-cell embedding *E*_cell_.

Formally, let *C* denote a single cell from the new dataset, *R* the original dataset’s consensus peak set, and *O* the set of overlapping regions between *C* and *R*. For each region *r*_*i*_ *∈ O*, we compute the embedding 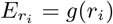, where *g* is the embedding transformation 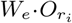. The whole-cell embedding is then computed by averaging these region embeddings:

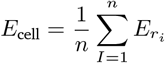

where *n* is the number of regions in *O*. These vector averages are computed with the `numpy` package in python.

#### EV-Projection

The EV-Projection procedure extends the E-Projection method by incorporating an additional visualization step. This approach retains all the benefits of E-Projection, while also providing a means to visualize the projection results using the pre-fit UMAP model derived from the original reference data embeddings.

As with E-Projection, we begin by calculating region overlaps and converting regions to their corresponding embeddings. We then average these embeddings to compute a whole-cell embedding for each cell in the new dataset.

After obtaining the whole-cell embeddings *E*_cell_ from E-Projection, these embeddings are further transformed using the UMAP model fitted to the original data embeddings. The explicit equation for UMAP’s transformation process on a new point is not straightforward to express in a simple mathematical formula due to the complexity of the algorithm [33]. To that end, we denote this transformation function as *f*_UMAP_. The EV-projection cell-embeddings as follows:

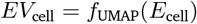

This transformation facilitates the visualization of the new data in the same low-dimensional space as the original data, providing a spatial relationship between new and original cells. This step aids in interpreting and comparing the accessibility landscape of the new cells in the context of the original cells.

### Cell type classification

#### Embeddings of the Luecken2021 dataset

To classify new, unseen single cells we first generate an embedding for every cell in the Luecken2021 dataset [25]. Each cell in the dataset, denoted as *x*_*i*_, is transformed into a highdimensional representation, or an embedding, denoted as *e*_*i*_ using the pre-trained model. Each embedding has an assigned ground-truth cell type label. The embeddings are generated with the no-projection procedure and stored alongside their metadata in a Qdrant database. Qdrant is open sourced and can be found on GitHub: https://github.com/qdrant/qdrant. Qdrant allows fast, convenient approximate nearest neighbor computation. We denote the set of all embeddings as *E* = *e*_1_, *e*_2_, …, *e*_*n*_, where *n* is the total number of cells in the Luecken2021 dataset. The corresponding cell type labels are represented as *L* = *l*_1_, *l*_2_, …, *l*_*n*_.

#### Approximate K-nearest-neighbor calculation

Given an unseen data point, *x*_*u*_, the goal is to assign it a label by using the embeddings of the pre-labeled dataset. The first step towards this is to compute an embedding for the unseen data point, through E-projection. Next, we compute the approximate nearest neighbors to *e*_*u*_ using navigable small world graphs with controllable hierarchy (Hierarchical NSW, HNSW) [38]. This is implemented in Qdrant with cosine distance and we simply query the database with the new embedding *e*_*u*_.

#### Label transfer

Once the approximate k-nearest-neighbors are retrieved, we assign a label to *e*_*u*_ by performing a consensus vote among its *k* nearest neighbors. We denote the indices of these *k* nearest neighbors as *I* = *i*_1_, *i*_2_, …, *i*_*k*_. We then assign a label, *l*_*u*_, to the unseen data point, *x*_*u*_, based on the most common label among its *k* nearest neighbors, which can be formalized as:

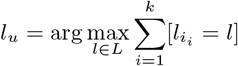

where 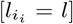 is the Iverson bracket notation that equals 1 if the condition inside the brackets is met, and equals 0 otherwise. We use the Python `collections. Counter` object from the standard library to perform these computations.

#### Cluster annotation

After performing the label transfer procedure, the newly clustered dataset, which we’ll denote as *C* = *C*_1_, *C*_2_, …, *C*_*m*_, where *m* is the total number of clusters and each cluster *C*_*j*_ is a set of data points, undergoes a final round of consensus voting for label assignment.

For each cluster *C*_*j*_, we count the frequency of each label amongst all the data points within the cluster. Let 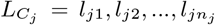 represent the labels of data points within cluster *C*_*j*_, where *n*_*j*_ is the total number of data points in *C*_*j*_ .

The label of cluster *C*_*j*_, denoted as 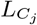, is then assigned based on the most frequent label among all data points in *C*_*j*_ . This can be formally defined as:

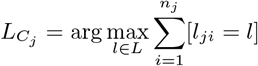

where [*l*_*ji*_ = *l*] is the Iverson bracket notation that equals 1 if the condition inside the brackets is met, and equals 0 otherwise.

In this way, each cluster is assigned the label that is most represented among its constituent data points. The quality of cluster labeling is highly dependent on the accuracy of the initial label transfer step.

### Evaluation of scEmbed cell type annotation

We use Cellcano, a novel scATAC-seq cell annotation method, to assign ground truth labels to our new PBMC data [12]. We follow the online tutorials (https://marvinquiet.github.io/Cellcano/) and leverage their provided reference dataset to process the new PBMC data. Once ground truth labels have been assigned by Cellcano and putative cell types are assigned from scEmbed, we can compute the F1 score to measure the accuracy of our classification.

The F1 score is the harmonic mean of precision and recall, and provides a balance between these two measures. Precision is the number of true positives divided by the sum of true positives and false positives, and recall is the number of true positives divided by the sum of true positives and false negatives. Formally, these are defined as:

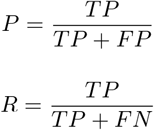

The F1 score is defined as:

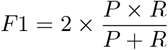

To compute these measures, we compare the predicted labels, denoted as *L*_*p*_ = *l*_*p*1_, *l*_*p*2_, …, *l*_*pn*_, to the ground truth labels, denoted as *L*_*g*_ = *l*_*g*1_, *l*_*g*2_, …, *l*_*gn*_, where *n* is the total number of data points (or clusters). We utilize the `metrics.f1 score` function from `scikit-learn` to compute this value.

## Data availability

scEmbed is open source and available at https://github.com/databio/geniml. Pre-trained models from this work can be obtained on hugging-face: https://huggingface.co/databio.

## Supplemental Information

**Supplementary Table S1.** Label mapping between scEmbed and cellcano for consistent comparison of classification performance.

| scEmbed label | Cellcano label |
| --- | --- |
| B1 B | B cells |
| CD4+ T activated | CD4 T cells |
| CD4+ T naive | CD4 T cells |
| CD8+ T | CD8 T cells |
| CD8+ T naive | CD8 T cells |
| CD14+ Mono | Monocytes |
| cDC2 | Dendritic cells |
| NK | NK cells |
| Naive CD20+ B | B cells |

**Supplementary Figure S1.**
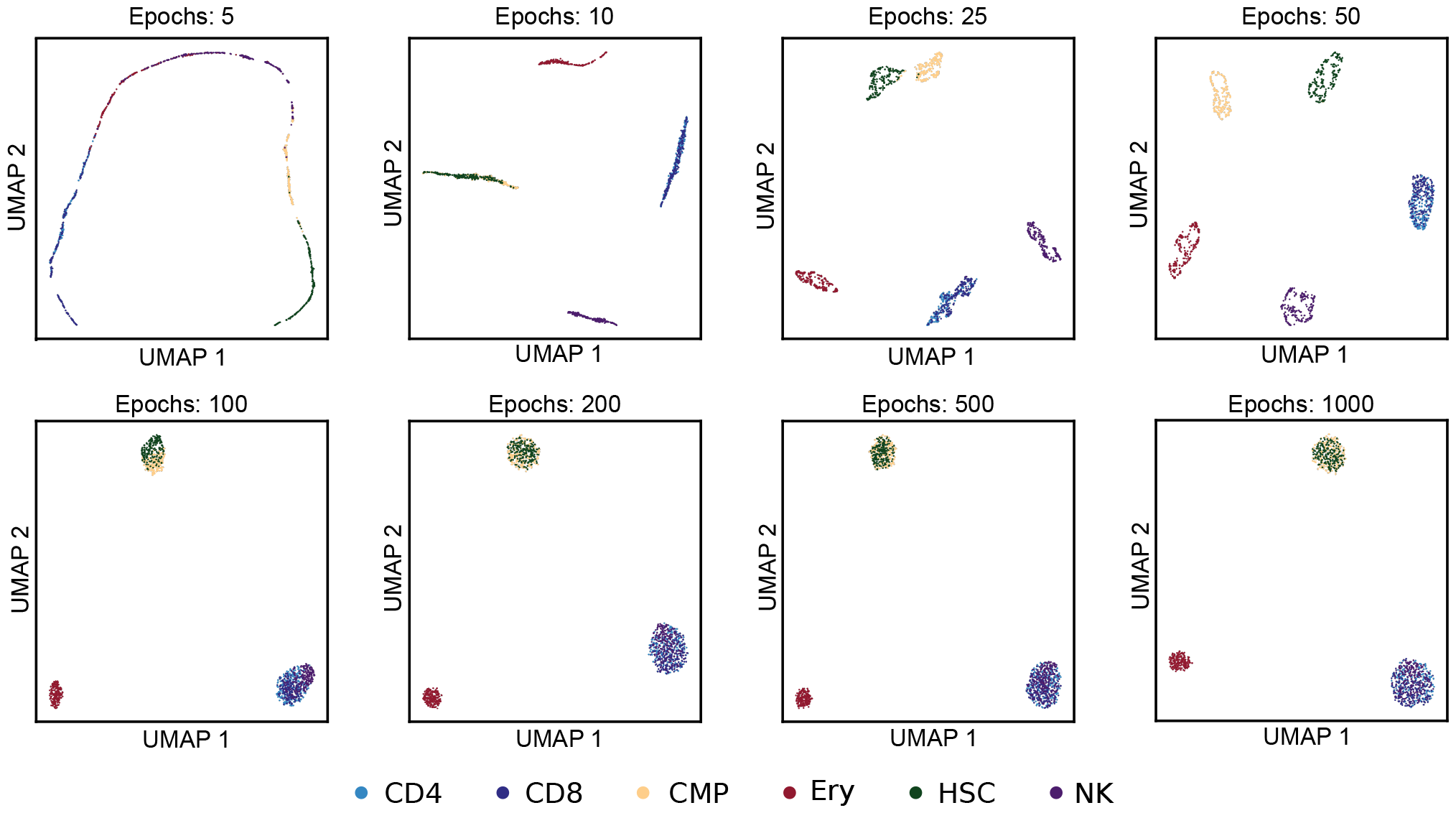
Epoch tests show that scEmbed learns well after 100 epochs. UMAP plots were generated after repeated model training using 8 different numbers of epochs for training. The model was trained on a synthetic bone marrow dataset described by Chen et al. UMAPs showed little change after 100 epochs.

**Supplementary Figure S2.**
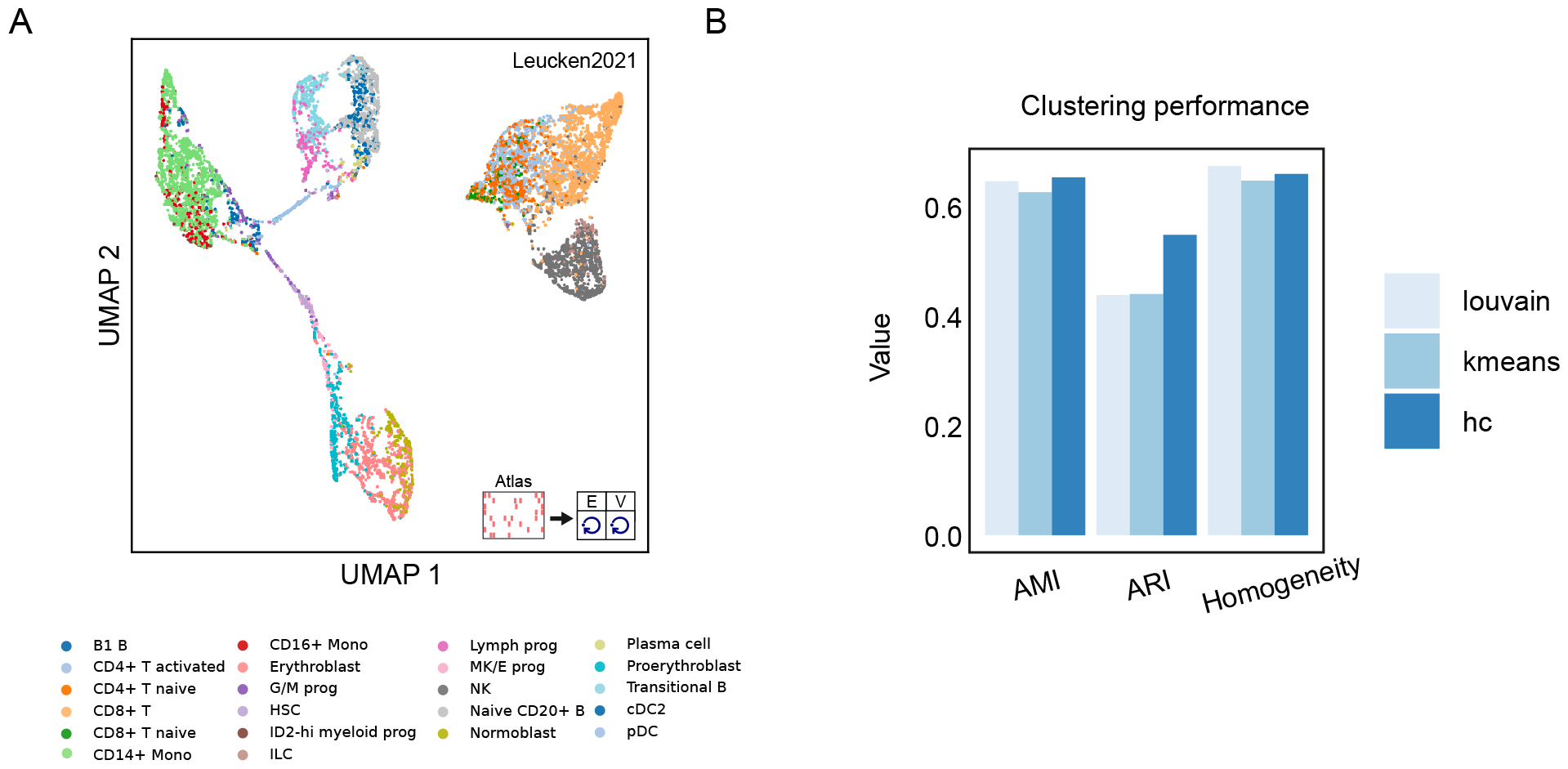
scEmbed produces visually distinct clusters of the Luecken2021 dataset. **A**. UMAP plot of the Luecken2021 dataset. **B**. ARI, AMI, and homogeneity scores for clusters produced by scEmbed using cell-type labels as the ground-truth.

**Supplementary Figure S3.**
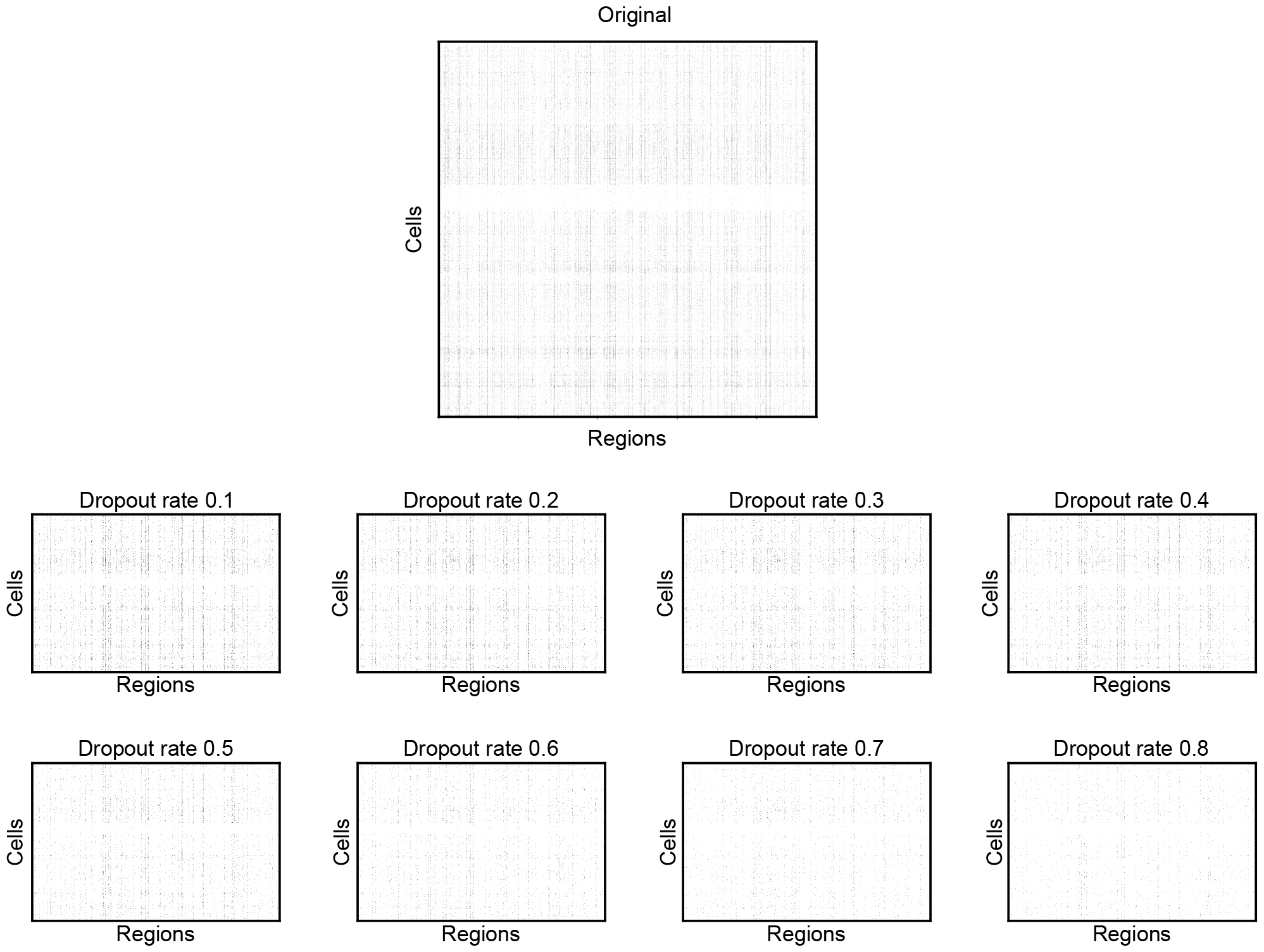
Sparsity plots enable visualization of iterative data dropout. Sparsity plots of the original matrix along with the matrix dropouts. Non-zero values decrease as the dropout rate increases from 10% to 80%.

**Supplementary Figure S4.**
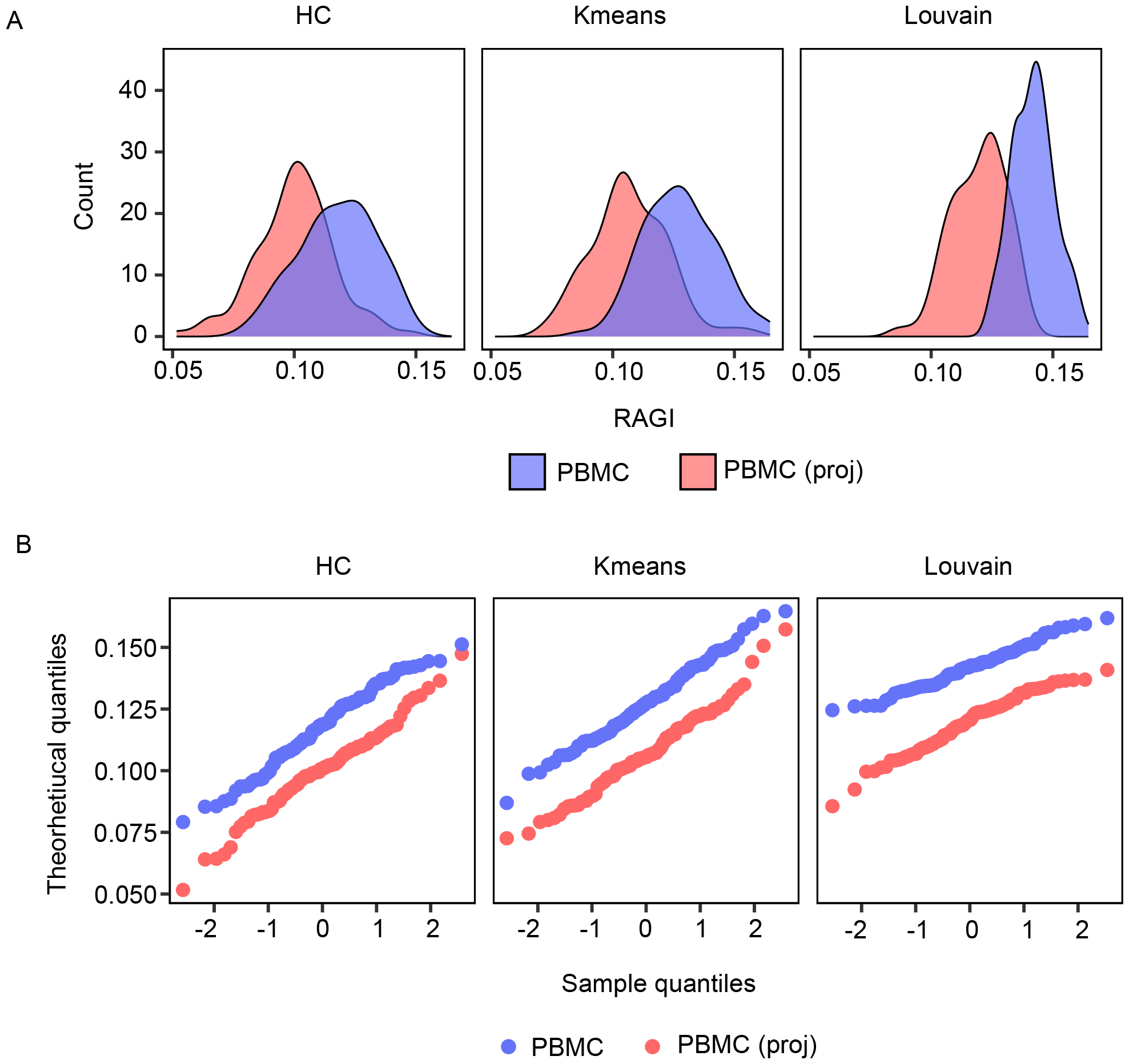
Distributions of the RAGI scores for all subsampled cells. **A**. Distribution of RAGI scores for cells with embeddings from the new model versus projection through the model trained on the Buenrostro2018 dataset. **B**. QQ plots of the RAGI scores for cells with embeddings from the new model versus projection through the model trained on the Buenrostro2018 dataset.

**Supplementary Figure S5.**
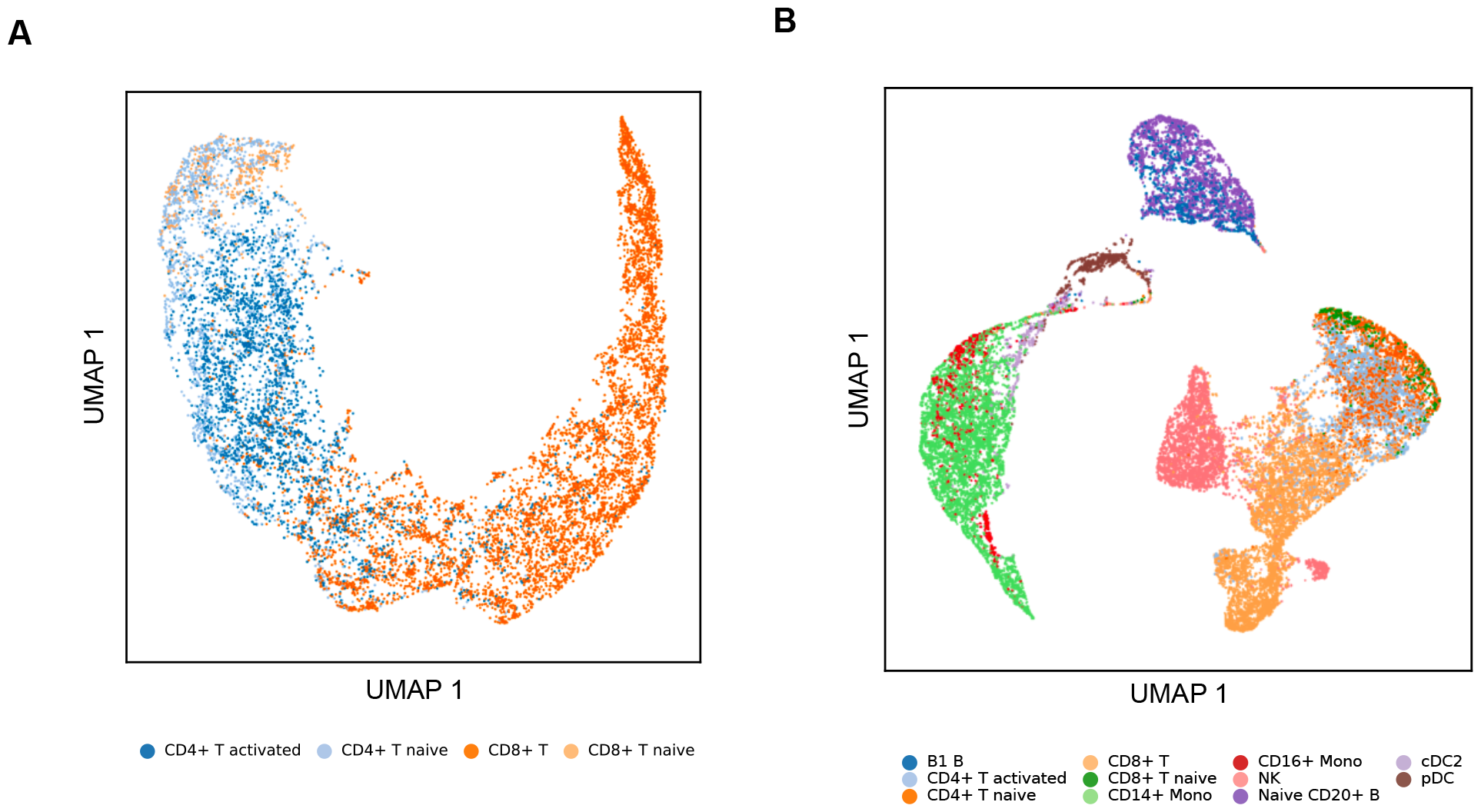
scEmbed produces visually distinct clusters of the subsetted Luecken2021 dataset. **A**. UMAP plot of an scEmbed model trained at 100 epochs for just the T Cells. Looking only at T Cells, the model can visually cluster the different cell types. B. UMAP plot of only PBMC cells from the Luecken2021 dataset.

**Supplementary Figure S6.**
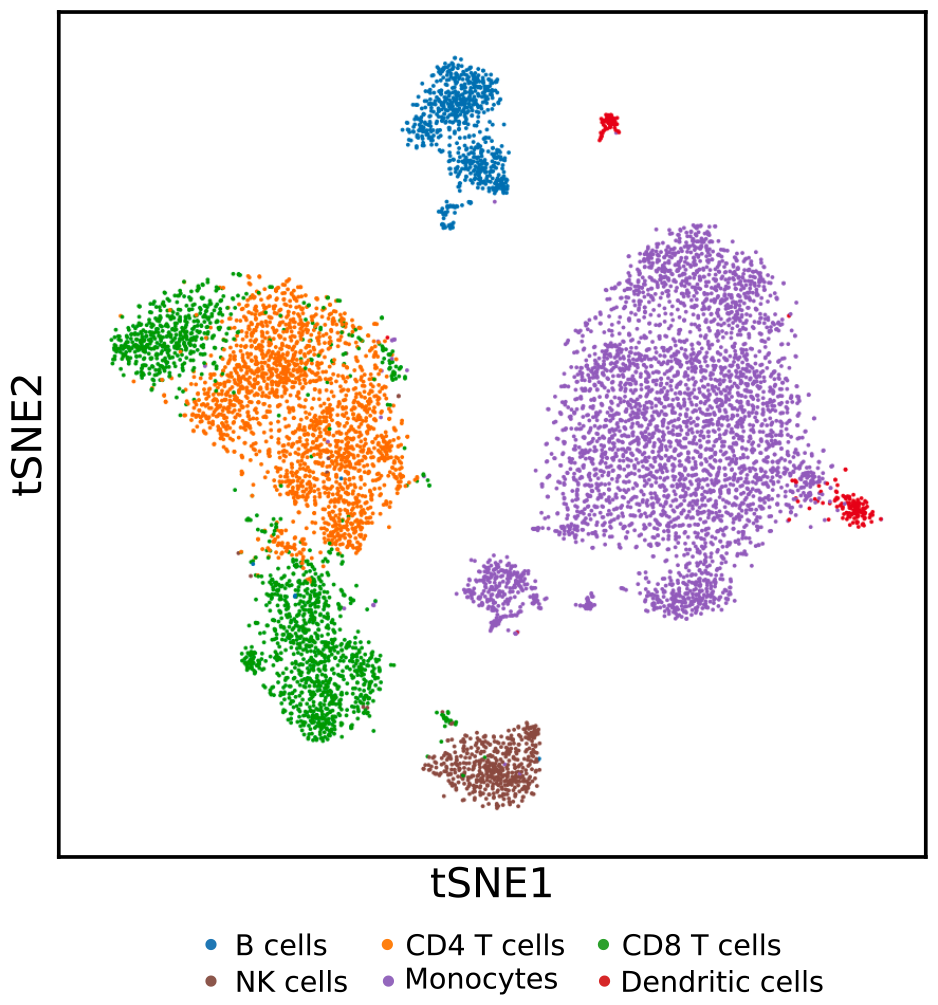
Visualization of reference data for Cellcano. tSNE plot output from cellcano showing clustered data from the given reference dataset.

**Supplementary Figure S7.**
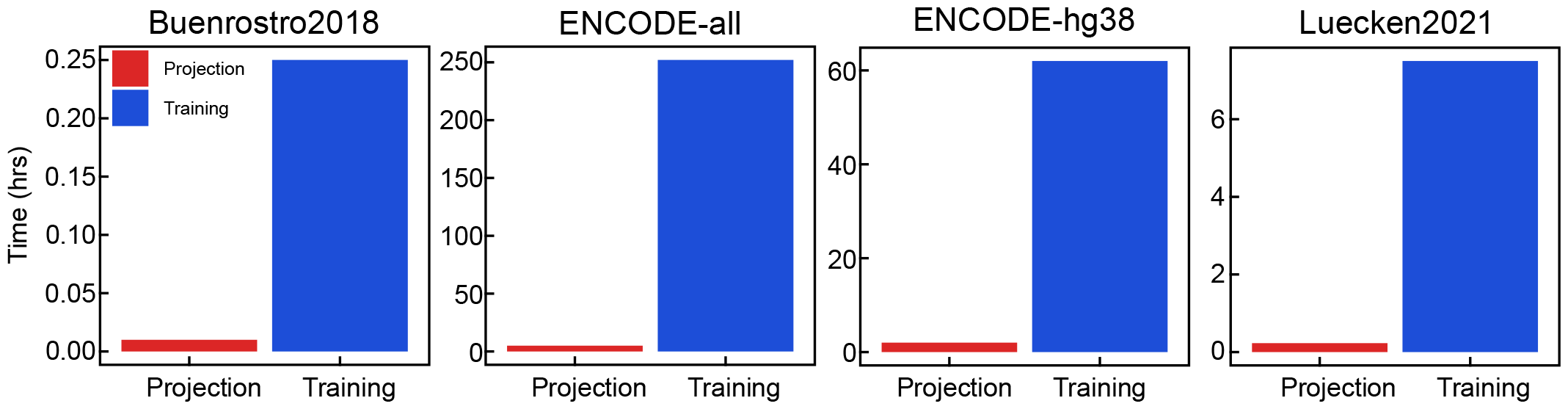
Number of hours required to perform training and projection for various datasets used in this study. Projection of new data into a pre-trained model takes very little time compared to training a model from scratch.

